# Given the birds, where is the flock? Visual estimation of the location of collections of points

**DOI:** 10.1101/2025.07.10.664170

**Authors:** Keiji Ota, Qihan Wu, Pascal Mamassian, Laurence T Maloney

## Abstract

Birds in flight form a flock, and pedestrians form a crowd. Humans readily perceive the center of such clouds, even though no individual bird or pedestrian marks that center. Developing accurate models of human perception of such phenomena is an important goal in vision research. Here, we reframed this central problem of perceptual organization in statistical terms: observers saw a sample drawn from an invisible probability density function (pdf) and attempted to estimate the location of that pdf, with better performance corresponding to lower estimation variance. Optimal minimum variance estimators differ across distribution families. We evaluated human performance in estimating the locations of pdfs from samples drawn from Gaussian, Laplacian, and Uniform location families. Observers approximated the family-specific optimal estimators for the Laplacian and Uniform distributions but deviated systematically from the optimum for the Gaussian distribution. How could the visual system flexibly adapt its estimates across different location families? We propose that observers adopt a single estimator based on clusters in the sample. According to this Visual Cluster Model, perceptual organization first aggregates individual ‘parts’ into clusters and only then combines these clusters into a representation of the ‘whole’, with the contribution of each cluster determined by the likelihood of the underlying distribution’s location. The model accounted for performance across all three distribution families, providing a general computational mechanism for estimating the location of a cloud from its parts across different statistical environments.

## Introduction

The human visual system segments scenes into visual objects (Koffka, 1935/2013; Wagemans, Elder, et al., 2012; Wagemans, Feldman, et al., 2012) adding visual structure to what are initially only patterns of light across the retinas. Such ‘wholes made of parts’ have their own spatial locations in 1D, 2D or 3D which the observer can be asked to estimate. If the observer is asked, for example, to saccade to a 2D cloud of points, the saccade can be interpreted as an estimate of the center of the cloud (McGowan et al., 1998; Melcher & Kowler, 1999). *The first question is, how is this location estimate computed from the individual points?* Birds in flight form a flock in 3D (Figure 1A) but typically no one bird marks the center of the flock. There is no obvious answer to the problem as posed.

**Figure 1:**
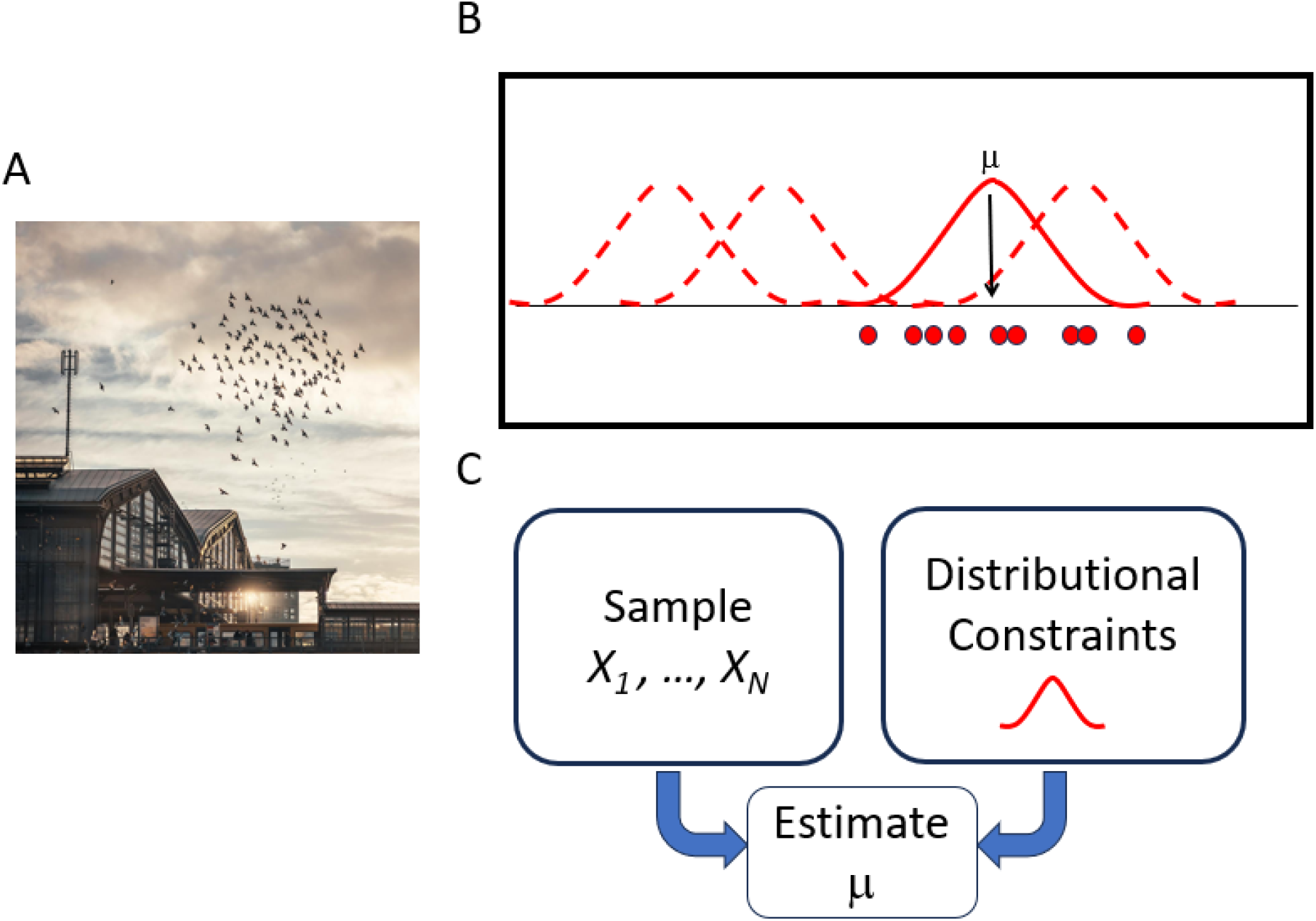
Estimating the Location of a Collection of Points. **A. A Flock of Birds.** A flock of birds in flight. Photo by George Frewat via Pexels, used under the Pexels license. **B. The one-dimensional Gaussian Location Task**. The dashed and solid red curves are examples of Gaussian probability density functions (pdfs) with unit variance for different values of the location parameter ***μ***. All of the red curves together comprise a *Gaussian location family*. The solid red curve is the actual pdf from which the sample was drawn. The sample is composed of *N* positions and is plotted as red points along the horizontal axis. The observer sees only the sample ***X***_**1**_, … ***X***_***N***_ and is asked to estimate the value of the location parameter ***μ*** corresponding to the pdf that was used to generate the sample. **C. 1D Parametric Estimation**. The observer sees the individual points and knows that the points are a sample from a pdf in the Gaussian location family with unit variance. He estimates the unknown location of the generating pdf and knowledge of the pdf may influence his choice of estimator.

We focus on a statistical version of the task where the ‘parts’ form a 1D sample drawn from a 1D statistical distribution and – importantly – we can compare human performance to normative. Different families of pdfs will serve as generators.

Not every perceptual task can be modeled as a statistical task where ‘parts’ are a sample drawn from a partially-known pdf, but considering such tasks allows us to use powerful statistical tools in a special case.

Other Gestalt phenomena such as border ownership (D’Angelo et al., 2025), object segmentation (Chen et al., 2025; Jeurissen et al., 2024; Papale et al., 2024) and shape perception (Nielsen & Connor, 2024) and more are also part of ‘middle vision’ (Ullman, 1984), the stage in visual processing that follows encoding of initial visual information into retinotopic areas and links local features to the items in the environment that generated them (Groen et al., 2017; Peirce, 2015).

### Terminology

The collection of points (‘*parts’*) forms a *sample* drawn froms a one-dimensional *probability density function* (pdf), the *generator*. In Figure 1B, the red curves, dashed and solid, are copies of the same pdf with the red, solid pdf being the pdf used to generate the sample. They represent a *location family*, a collection of pdfs differing only in location (e.g. Figure 1B). The colored dots are a 1D sample from the solid pdf. The location estimation problem is a problem in parametric point estimation (Lehmann, 1998; Mood et al., 1974) as summarized in Figure 1C.

We address three questions: (1) *How does the observer estimate the location of the pdf given only the sample and knowledge of the location family to which the pdf belongs?* (2) *How does changing the pdf (‘generator’) underlying the location family affect the location estimation – if it has any effect at all? And* (3) *How does human location estimation compare to uniform minimum variance unbiased estimation (UMVUE) for each of the distributions we consider?*

The three location families we consider are the Gaussian, the Laplacian and the Uniform, illustrated below (Figure 2). A possible outcome is that the observer uses the same estimator, possibly the Gaussian, for all three distributions. A second possibility is that the observer uses different estimators for the different location distributions but at least one of them is not the UMVUE of the corresponding distribution. A final possibility is that the observer chooses estimators close to the UMVUE for each of the three location distributions, somehow adapting its estimation algorithm to the distribution (generator).

**Figure 2.**
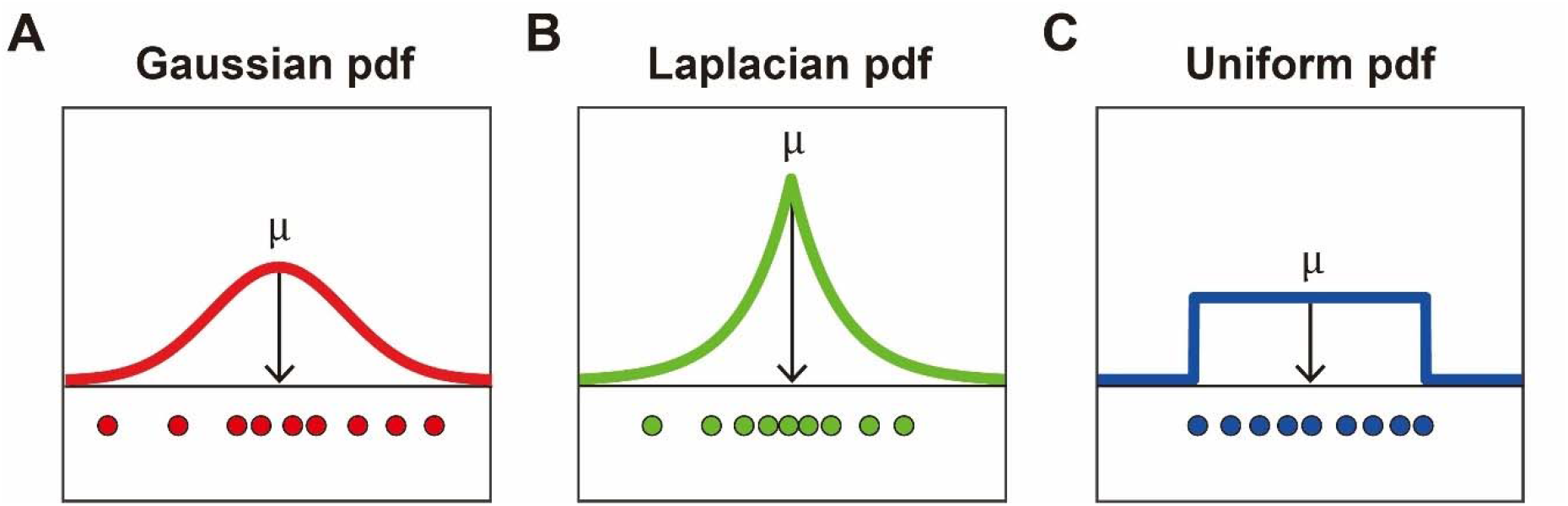
Location Tasks. **A. Gaussian Location Task. B. Laplacian Location Task. C. Uniform Location Task.** Each plot shows a probability function and a sample, color coded. The pdfs used in the experiment all had the same variance. See Materials and Methods.

In the present study, we consider three different location-scale families of probability density functions (pdfs). One of these families is the Gaussian pdf, commonly used in visual modeling, notably cue combination models (Landy et al., 2007; Oruç et al., 2003). A second motivation for using Gaussian distributions in modeling visual judgments is that actual visual and motor error is close to Gaussian in a wide class of everyday tasks (Ota et al., 2019; Tanae et al., 2021; Trommershauser et al., 2008). Humans frequently encounter tasks involving Gaussian random variables and they may tend to apply estimation rules and other statistical procedures that are appropriate when distributions are Gaussian (Tassinari et al., 2006; Vilares et al., 2012). However, it is less well known how humans behave when distributions are non-Gaussian (see (Maloney & Thomas, 1991) for an exception).

Figure 1C summarizes what the observer in our experiment knows – the locations of points and knowledge of the distributional family from which they are drawn but not the precise pdf. *Parametric point estimation* methods (Lehmann, 1998; Mood et al., 1974) allow us to compute optimal estimators for each pdf family based on criteria such as minimum variance for a wide range of distributions.

If the pdf is Gaussian with fixed variance 1 and with unknown *location* parameter ***μ***, the pdf is

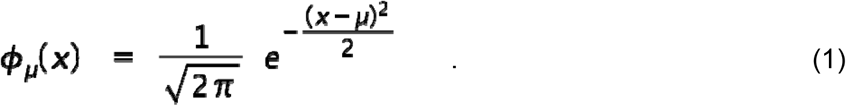

Several examples of pdfs from a *Gaussian location family* (Eq. 1) are plotted as dashed red curves in Figure 1B. As ***μ*** is varied, the pdf moves left and right (hence the term “location parameter”). The “true” pdf, the one from which the sample was actually drawn, is plotted as a solid red curve.

Again, the observer’s task is – given only the sample and the Gaussian constraint – to estimate the unknown *location* ***μ*** of the true pdf. The estimator that the observer uses is a function of the sample and may depend as well on the distributional assumption (Figure 1C). The observer’s estimate can be denoted 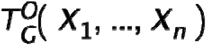 or, if there is no ambiguity, just 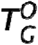. “O” stands for the Observer, “G” for the Gaussian assumption.

While the observer has two sources of information (*distributional constraints* and the *sample*) that inform their estimate 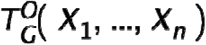, it is usually the sample values that take center stage, with the Gaussian assumption in the background. But what use – if any – does the observer make of the distributional constraints? How are the parts combined to estimate the location of the whole when the distribution generating the parts is not Gaussian?

In Figure 2 we illustrate the Location Task for three choices of distributional family. Figure 2A corresponds to the Gaussian case that we just discussed (Eq. 1). Figure 2B illustrates a location task based on the Laplacian distribution^1^, where the pdf is now

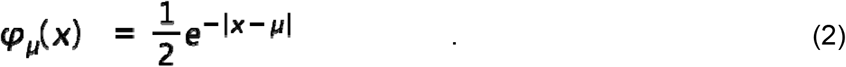

The pdf is symmetric around the unknown location parameter ***μ***. The observer’s task is still to estimate ***μ***. Finally, the Uniform Location Task is based on a uniform distribution (Figure 2C),

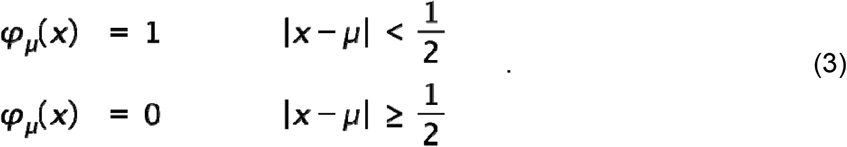

We assessed human performance in the three location tasks for the distributions shown in Figure 2. In our experiment, the observer familiarized himself with the distributional families and the typical samples they generate. On every training trial, they saw a sample drawn from a pdf. The color of the points identified the pdf location family from which the pdf is drawn (as in Figure 2).

### Uniformly Minimum Variance Unbiased Estimators (UMVUE)

An estimator is any function of the sample, ***T***(***X***_**1**_, … ***X***_***n***_) and, since an estimator is a function of random variables, it is itself a random variable with its own expected value ***E*[*t*]** and variance ***Var*** (***T***).

A common criterion for a “good” estimator is that it is *unbiased*, i.e. ***E*[*t*] = *μ***. A second common criterion is that the estimator should have the lowest possible variance. If an estimator satisfies both of these two criteria for all values of ***μ*** then it is a *Uniformly*

*Minimum Variance Unbiased Estimator (UMVUE)* or, less formally, the *minimum variance estimator*. Cue combination models use UMVUEs and compare these with estimators human observers adopt (Landy et al., 1995; Oruç et al., 2003; Trommershauser et al., 2011).

We can compute a measure of the *efficiency* of any unbiased estimator ***T*** for a task as the ratio of the variance of the UMVUE to that of the estimator. Since 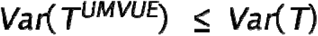, the *efficiency* of ***T***

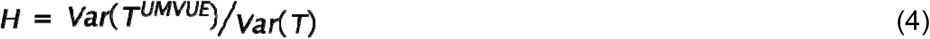

is a value between 0 and 1.

#### L-Estimators

For convenience, we will focus on a set of estimators referred to as L-estimators (Lehmann, 1998). They are defined as weighted combinations of order statistics

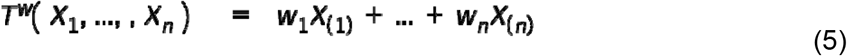

where the sum of the weights is constrained to be 1. The order statistics (Mood et al., 1974) are just the sample values renumbered from left to right. We can represent any L-estimator by a plot of its weights (Figure 3).

**Figure 3:**
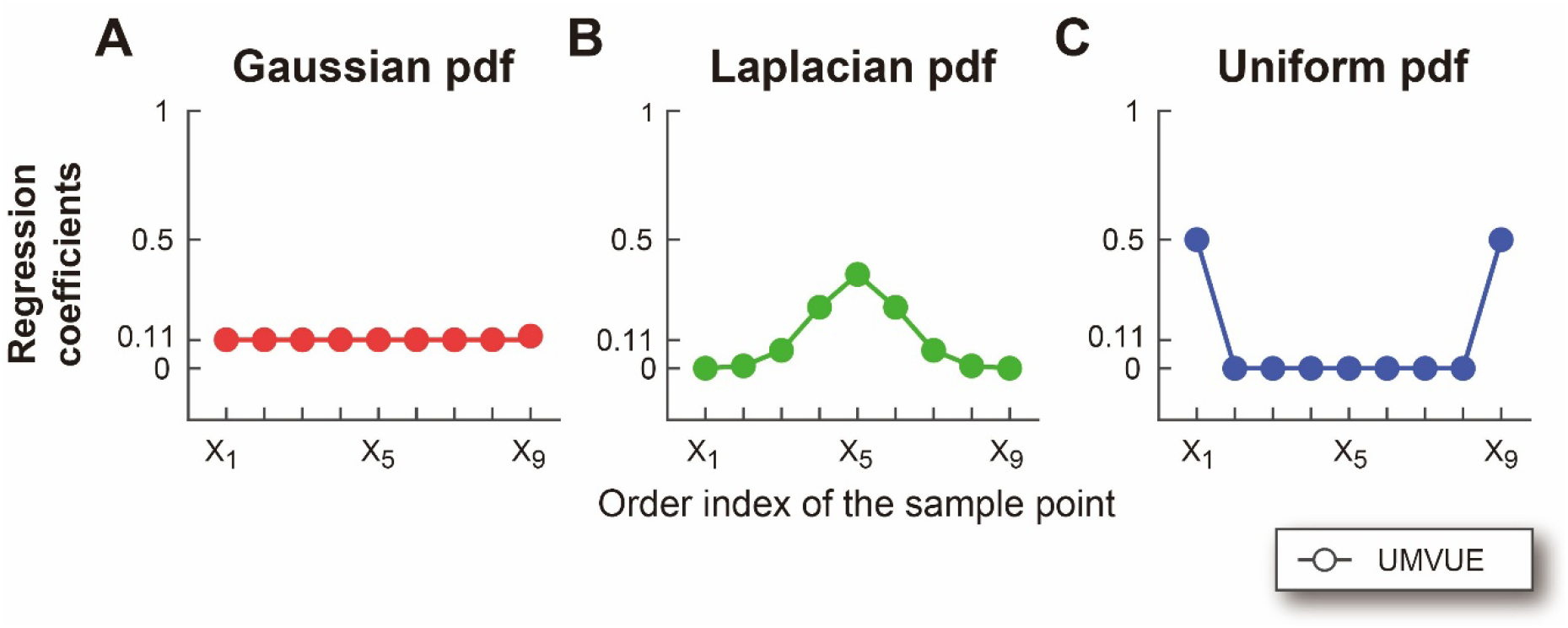
Uniformly Minimum Variance Unbiased Estimators (UMVUEs). **A. Gaussian Location Task. B. Laplacian Location Task.** The *UMVUE*_*L*_ L-estimator estimated numerically (the Laplacian has no closed-form solution for its *UMVUE*). **C. Uniform Location Task**. Three UMVUE estimators are represented as L-estimators of weighted average of order statistics. The Y-axis denotes weights (i.e., regression coefficients) on each sample point ordered from left to right. UMVUE estimator for the Gaussian location task is the arithmetic mean of the sample. The UMVUE for the Laplacian location task has the largest weight on the median. The UMVUE for the uniform location task is the average of extremes.

### The UMVUEs for the Three Location Tasks

Parametric estimation comprises a collection of mathematical techniques that allow us to identify optimal estimators including UMVUEs for different estimation problems with different distributional families (Lehmann, 1998; Mood et al., 1974). The unbiased estimator ***T***^**+**^ minimizing variance for the Gaussian task illustrated in Figure 1B is just the arithmetic mean of the sample (Lehmann, 1998; Mood et al., 1974). It is repeated in Eq. 6, written in the form of an L-estimator

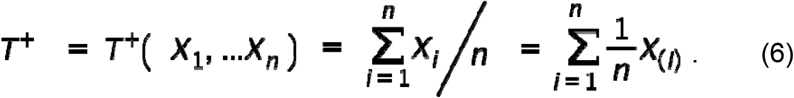

This estimator is unbiased and it minimizes the variance ***E*[*t***^**+**^ **− *μ*)**^**2**^].

Not every location task has an *UMVUE*. While the *UMVUE* for the Gaussian is the sample *mean* (Eq. 6), the Laplacian has no closed-form solution for its *UMVUE*. We computed the unbiased L-estimator that minimized variance by numerical methods (exhaustive search) and plotted it in Figure 3B, the *UMVUE* among L-estimators denoted *UMVUE*_*L*_. We use the term *UMVUE* to refer to either *UMVUE* or *UMVUE*_*L*_ when it is clear from context which is intended.

The *UMVUE* for the uniform family is

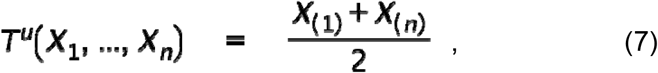

the average of the minimum and maximum values in the sample (the *average of extremes*). We have three different *UMVUEs* each matched to the distribution family of a location estimation task (Fig 3). Each is unbiased.

Previous studies on human vision have typically used the Gaussian location family in modeling human performance. These studies have shown that human observers accurately estimate the center of a 2D cloud of points and thus adopt the correct UMVUE for a location estimation of a Gaussian distribution (Gan et al., 2023; Juni et al., 2010; McGowan et al., 1998; Melcher & Kowler, 1999; Rashid & Chubb, 2022; Sun et al., 2016, 2018). However, the centroid is not the minimum variance estimator for the locations of Laplacian and uniform distributions. It has not been tested how a different location family of a statistical distribution systematically influences human location estimation based on the individual points.

We will investigate how the human visual system adapts its estimation rule *T*^*o*^ (*X*_1_, … *X*_*n*_) to location tasks based on each of the distributional families in Figure 2 – if it adapts it at all? Second, we will compare human performance to optimal unbiased estimators minimizing variance (UMVUEs) for each of the three location families in Figure 2.

One possible outcome is that the observer may simply adopt the same estimator for all three location tasks, ignoring the distributional information. At the other extreme, the observer might adopt the correct UMVUE estimator for each of the three tasks in Figure 3.

#### Influence

We can interpret the weights in an L-estimator as measures of the *influence* of each point (each order statistic), a useful characterization of a statistical task (Juni et al., 2010; Maloney & Landy, 1989; Ota & Maloney, 2024) that is important for research in robust statistical estimation (Hampel et al., 1986; Huber, 1981).

In our experiment, we are presenting the observer with a sample whose order statistics — left to right – are and ***X***_**(1)**_, … ***X***_**(*I*)**_, …, ***X***_**(*n*)**_ recording the observer’s response ***T***^**o**^**(*X***_**(1)**_, … ***X***_**(*I*)**_, …, ***X***_**(*n*)**_), the location estimate. The influence of the point at position *i* on the location estimate is the partial derivative

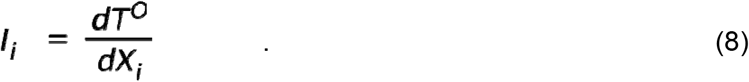

which allows us to measure the contribution of each point to the overall estimate. If the location estimator is an L-estimator, then the influence estimates are just the weights in Eq. (5). We compare the normative influence measures for each task (Figure 3) with the observed influence measures computed by linear regression.

To summarize, in this article we first examine how observers combine locations of individual points to estimate the location of the generating pdf, an exemplary parametric estimation problem in visual form. We then propose a model of human performance, the Visual Cluster Model (VCM) of location estimation. We test the VCM model against alternatives.

## Results

The observers were first familiarized with the three distributional families (Gaussian, Laplacian, and Uniform) by viewing the typical samples generated for different sample sizes (Supplementary Figure 1A; see Materials and Methods). In the main experiment observers were presented with samples of 9 points drawn from one of the three distributional families and asked to estimate the position of the center of the generating pdf given only 9 sample points and the knowledge of which distribution family had generated the sample. They were given feedback concerning their visual estimate (Supplementary Figure 1B). We confirmed that behavioral performance did not systematically change across trials (Supplementary Figure 2A and 2B).

We know the UMVUE weights (influences) for each task (Figure 3) and we can estimate the observers’ weights by linear regression (see Materials and Methods). Figure 4 (and Supplementary Figure 3A) illustrates the estimated weights for each participant and the average across participants. Observers were sensitive to the distributional constraints. Observers’ influence measure markedly changed with changes in distributional family (interaction between the order of points and the distributional families by ANOVA on the estimated weights: *F* [16, 304] = 44.53, *p* < 0.0001, partial *η*^2^ = 0.701).

**Figure 4:**
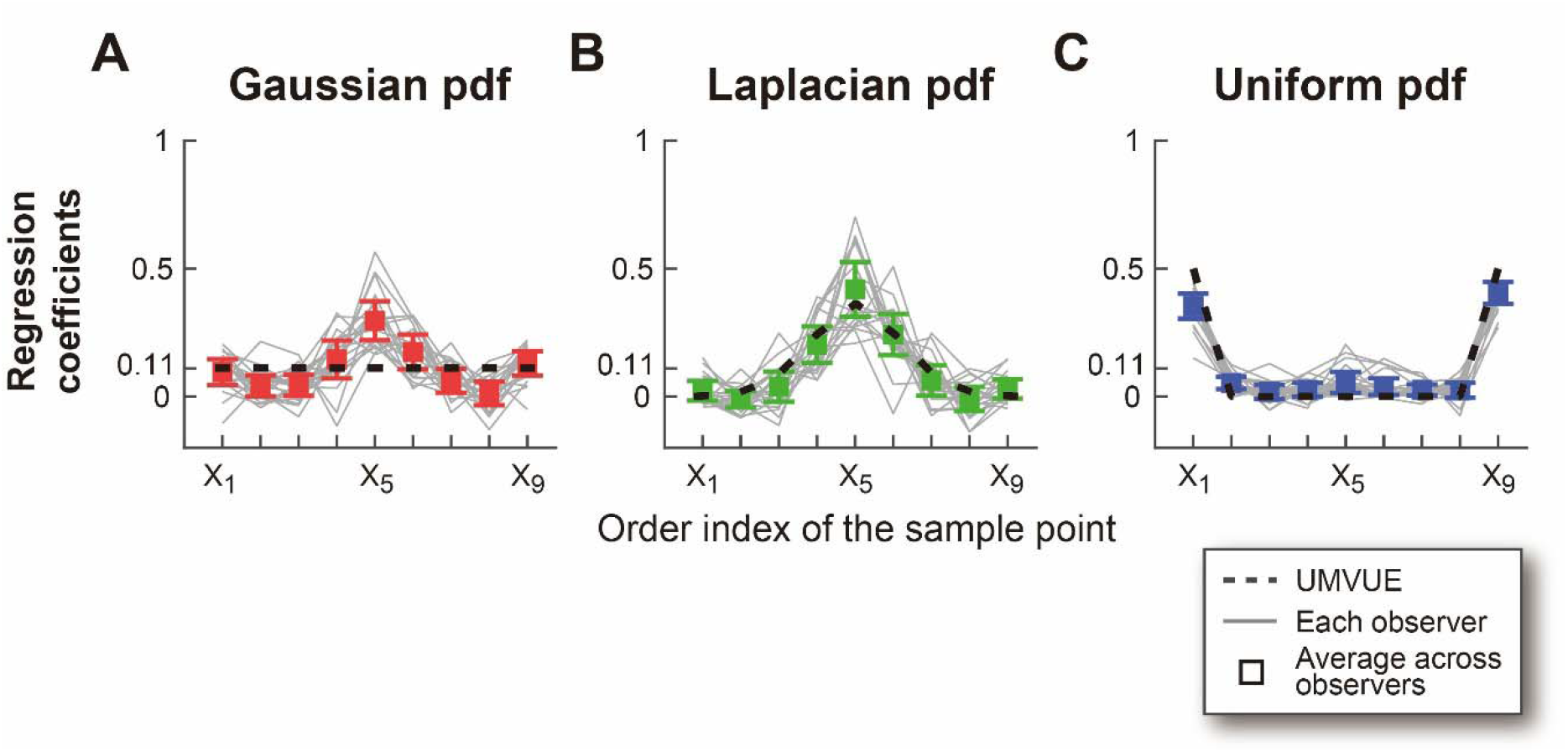
Observers’ Estimated Influence Measures Compared to UMVUE Influence Measures. **A. Gaussian Location Task. B. Laplacian Location Task. C. Uniform Location Task.** Influence (i.e. regression coefficients) of each point in samples (ordered from left to right) on observers’ visual estimates. Results for each observer are plotted as faint grey lines. Mean results across observers are plotted as color-coded squares. **The** error bars denote 99.7% confidence interval (± 3 SEM). Weights employed by UMVUE are taken from Figure 3 and replotted as dashed lines. For the Laplacian distribution, observers’ estimated influence measures are not significantly different from the UMVUEL. The observers’ influence measures for the other two distributions deviate from those of their respective UMVUEs.

Do observers’ choices of estimators approximate the UMVUEs for each location task? We compared measured influence with normative (UMVUE or UMVUE_L_) influence for each location task separately. For the Gaussian task, the normative estimator is just the arithmetic mean of the sample. In contrast, the observers’ estimated influences look W-shaped and are significantly different from the normative influences (*F* [8, 152] = 19.56, *p* < 0.0001, partial *η*^2^ = 0.507) by particularly placing larger weights on the median sample point than UMVUE (Supplementary Figure 3B), meaning that observers do something other than averaging points.

For the Uniform task, the measured influences are also different from normative influences (*F* [8, 152] = 35.27, *p* < 0.0001, partial *η*^2^ = 0.650). Even though observers gave most weight to the extreme sample points, they placed significantly less weight on them than the UMVUE (Supplementary Figure 3B). For the Laplacian task, observers approximated the UMVUE (*F* [8, 152] = 1.83, *p* = 0.076, partial *η*^2^ = 0.088). We note that there were no significant changes in the measured influences from the beginning to the end of the experiment (Supplementary Figure 2C). The efficiency of the observers’ estimators (Eq. 4) was overall 69.6% (Supplementary Figure 4). In other words, the observers were 1.44 times more variable than the UMVUE. There was no significant difference across the three location tasks (the main effect of the distributional families by ANOVA on the efficiency: *F* [2, 38] = 2.53, *p* = 0.093, partial *η*^2^ = 0.118).

In summary, the observers are sensitive to the distributional constraints in the three location tasks – choosing significantly different estimators for the three location tasks. The observers’ estimators did not differ significantly from the UMVUE_L_ for the Laplacian location task and roughly approximated the UMVUE for the Uniform location task.

### The Visual Cluster Model (VCM)

The observers used different estimators for the three location families. *Here, we considered the possibility that the visual system’s choice of estimator is based entirely on the cluster structure of the samples (Sun et al*., *2019)*. Intuitively, clusters convey information about the pdfs from which they are drawn. A cluster confined to an interval [a,b] will tend to have more points when the probability density across the interval is larger. We first illustrate the computation with an example.

In Figure 5A we plot a 9-point sample and in Figure 5B we partition the sample into *K* = 5 non-overlapping clusters denoted left to right as *C*_*1*_, *C*_*2*_, …, *C*_*5*_ using a standard clustering algorithm described below. Clusters in our experiment have a minimum size of 1 element, and a maximum size of 9 (in which case, there is only one cluster). We assign a location *c*_*j*_ to the *j*^th^ cluster *C*_*j*_, the midpoint of the endpoints of the cluster.

**Figure 5:**
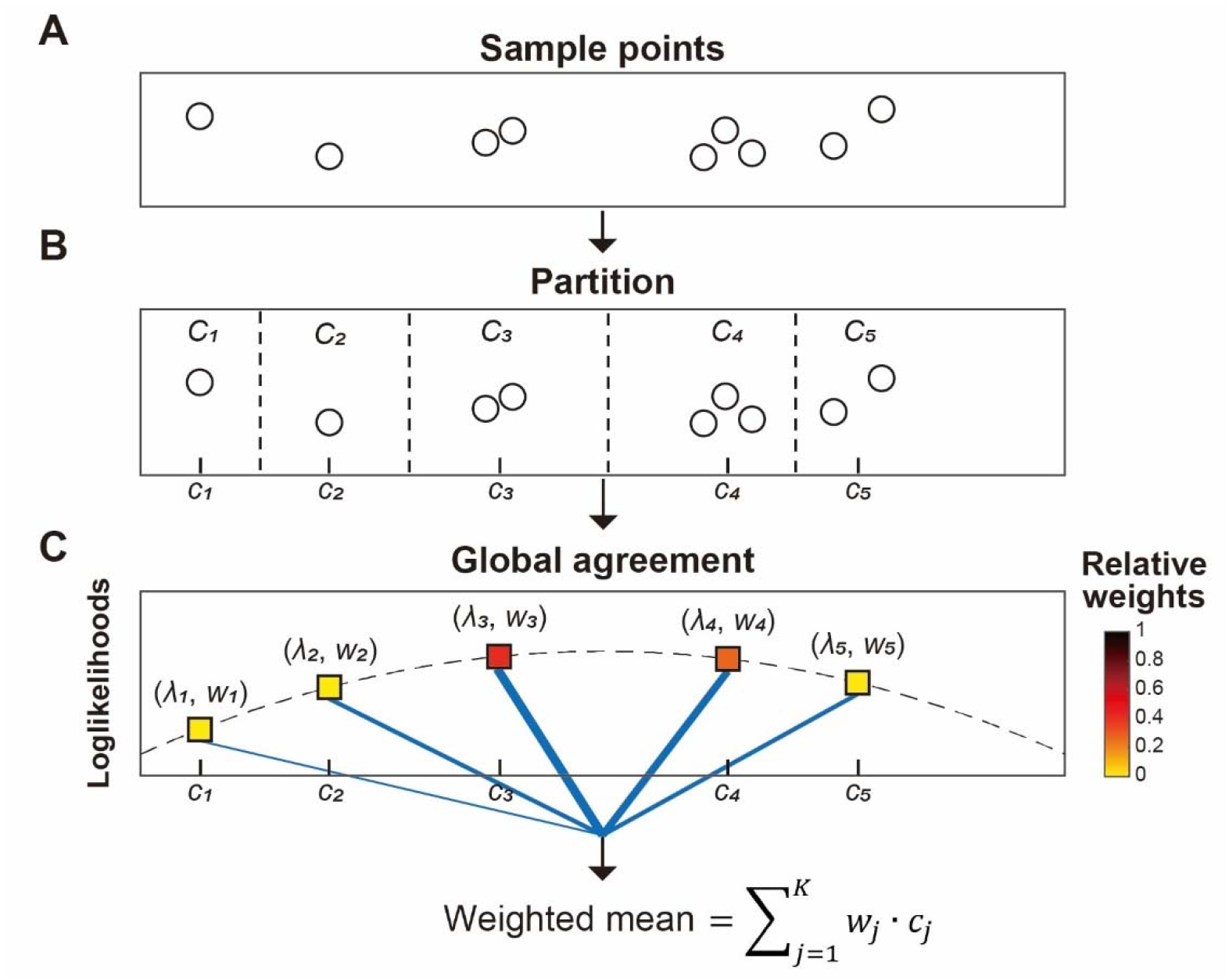
Visual Cluster Model. **A. Gaussian Location task.** A sample of 9 points is drawn from a Gaussian distribution. Observers were asked to estimate the location of the invisible center of the underlying distribution given only sample points. **B. Partition**. Visual cluster model partitions points (partitions shown as dashed vertical lines) into *K* clusters of the sample, ***C***_**1**_, … ***C***_***K***_. Each cluster is assigned a location ***C***_**1**_, … ***C***_***K***_ defined as the average of its largest and smallest points if the cluster contains more than one point. Otherwise, the location of the single point is the location of the cluster. **C. Global Agreement**. Visual cluster model assigns the log-likelihood *λ* _*j*_ of the center of the pdf to the location *c* _*j*_ of the *j*^*th*^ cluster. A dashed curve represents the log-likelihood values across the entire space of the task. The log-likelihood values *λ*_1_,…,*λ*_*K*_ are converted to the relative weights *w*_1_,…,*w*_*K*_ via a softmax transformation. The visual cluster model estimates the position of the center of the pdf as a weighted average of cluster locations *c*_1_,…,*c*_*K*_ with weights *w*_1_,…,*w*_*K*_.

In our example, the largest cluster, *C*_*4*_, contains three points. If we had to base our estimate of location on just one of the clusters in the cluster structure of the sample, it would plausibly be the 4th cluster. But there are competing clusters with either 1 or 2 points, and we can attempt to improve our estimator by weighting together the locations of the clusters with weights that depend on the clusters.

We considered variations on the model at each step but – to simplify the presentation of the model – we first describe one version of the model and discuss possible variations only after presentation of this base version of the model. One goal of the model is to reduce the complexity of the computation (Lieder & Griffiths, 2019) by relying on the cluster structure of the model (five clusters in our example) (Sun et al., 2019) rather than the nine points.

#### Step 1: Partition

The visual system partitions points in the sample into a small set of non-overlapping clusters. In Figures 5A and 5B, we apply a standard clustering algorithm, K-means clustering (Hartigan & Wong, 1979), to a sample ***X***_**1**_, … ***X***_***N***_ from the Gaussian distribution (Eq. 1). The K-means clustering algorithm has a free parameter *K* (the number of clusters). In Figure 5B, we obtained *K* = 5. The result is a partition into non-overlapping clusters numbered from left to right, possibly varying in number of points. We assign a location to each cluster ***C***_**1**_, … ***C***_***K***_ as the average of its largest and smallest points (average of extremes) if the cluster contains more than one point, otherwise the location of the single point if the cluster contains only one. The first step is analogous to the cluster-of-samples representation developed by Sun et al. (2019). We extend their model by introducing *global agreement*.

#### Step 2: Global agreement

How well does each cluster *C*_*j*_ at location *c*_*j*_ capture not just the points in its own cluster but rather the entire sample of points ***X***_**1**_, … ***X***_***N***_? We hypothesize that observers make use of the distributional constraint to assess the likelihood of the location of the distribution generating the points. In Figure 5C, for each cluster, we compute the log-likelihood *λ* _*j*_ of the location parameter ***μ*** of the population pdf with respect to the location *c* of the *j* ^th^ cluster:

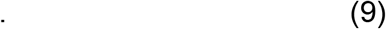

where *f*_*μ*_ is the likelihood function of the distribution chosen to generate the sample points, with unit variance. This step results in a measure of the likelihood that the location *c*_*j*_ of cluster *j* reflects the center of the distribution, taking into account all the points of the entire sample. Therefore, we refer to it as *global agreement*. A dashed curve in Figure 5C shows the log-likelihood computed across the entire space of the task. In the Gaussian location task, finding the location of the maximum log-likelihood value will correspond to averaging sample points. Instead of performing the log-likelihood computation over the entire space, we reduced the complexity of computation (Lieder & Griffiths, 2019) by computing the log-likelihood value with respect to the locations *c* of the *j* ^th^ clusters.

We use a softmax transformation with one parameter ***β***, i.e., known as inverse temperature (Reverdy & Leonard, 2016) to convert the log-likelihood *λ* _*j*_ to weights

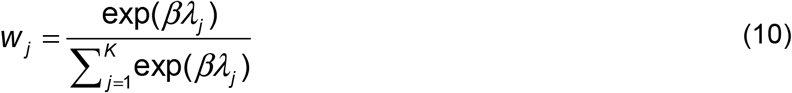

The parameter ***β*** is a free parameter used to scale measures of global agreement. When ***β*** is fixed at 0, the relative weights *w*_*j*_ are all 1/K: all clusters get equal weight. When ***β*** increases, clusters having a large log-likelihood value get relatively larger weights relative to those with smaller log-likelihood values. In the limit as grows, the weight of the cluster with the largest log-likelihood converges to 1 while the other weights drop to 0.

On any given trial, the visual system’s estimate of the position of the center of the distribution is modelled as the weighted mean of the locations of the clusters with softmax weights that depend on the log-likelihoods assigned to clusters (Figure 5C)

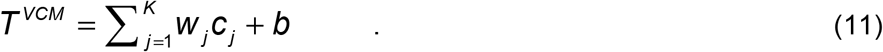

A bias term ***b*** is added to account for the small but constant bias that we found in the participants’ visual estimates (on average, 2.07 mm, SD = 1.41 mm), possibly a visuo-motor error.

The weight on each cluster is determined by the value of the measure of global agreement (i.e., how accurately the center of the distribution is approximated) relative to the measures of global agreement in other clusters.

#### Factorial model comparison

We test two levels of the grouping parameter *K*. At the first level, we will fix *K* at 9. At the second level, the grouping parameter *K* will be treated as a free parameter and estimated separately from the data in each location task (maximum likelihood). Given that samples from the three possible distributions are color-coded, the observer can select different values *K*_*G*_,*K*_*L*_, *K*_*U*_ of the grouping parameter *K* for the three different distributional families.

For global agreement (***β***) we test three levels in the factorial design. At the first level, we will fix ***β*** at 0 so all clusters get equal weights. At the second level, the parameter ***β*** will be treated as a free parameter constrained to be the same value across the three distributional families. Last, at the third level, we will fix it at 1. By this constraint, global agreement is applied to the model fit but the ***β*** value is not informed from each observer’s data.

We built six models by combining two levels of partition and three levels of global agreement and compare the models by a factorial model comparison (van den Berg et al., 2014; Zhang et al., 2020). Figure 6A shows the summed AICc score across participants for the six models. The two-stage visual cluster model (partition and global agreement) outperformed the other five variants of the model. A group-level Bayesian model selection (Rigoux et al., 2014; Stephan et al., 2009) showed that the protected exceedance probability of the two-stage VCM, i.e. the probability for the model to outperform all the other models was 99.9%. This model was also better than the normative model where UMVUE was fitted to the observer’s data. Furthermore, the three models based on partitioning points (*K*_*G*_, *K*_*L*_, *K*_*U*_ estimated from the data) outperformed the other three models without partitioning (*K* fixed at 9; the protected exceedance probability = 99.8%). A model where partitioning was applied while global agreement was partially applied (***β β*** fixed at 1) outperformed a model where partitioning was applied but global agreement was not (***β*** fixed at 0), though the ***β*** value was not fitted to the data in these two models (the protected exceedance probability = 87.1%). *These results support the claim that both partitioning and global agreement are essential computations that underpin the observers’ visual judgments*.

**Figure 6:**
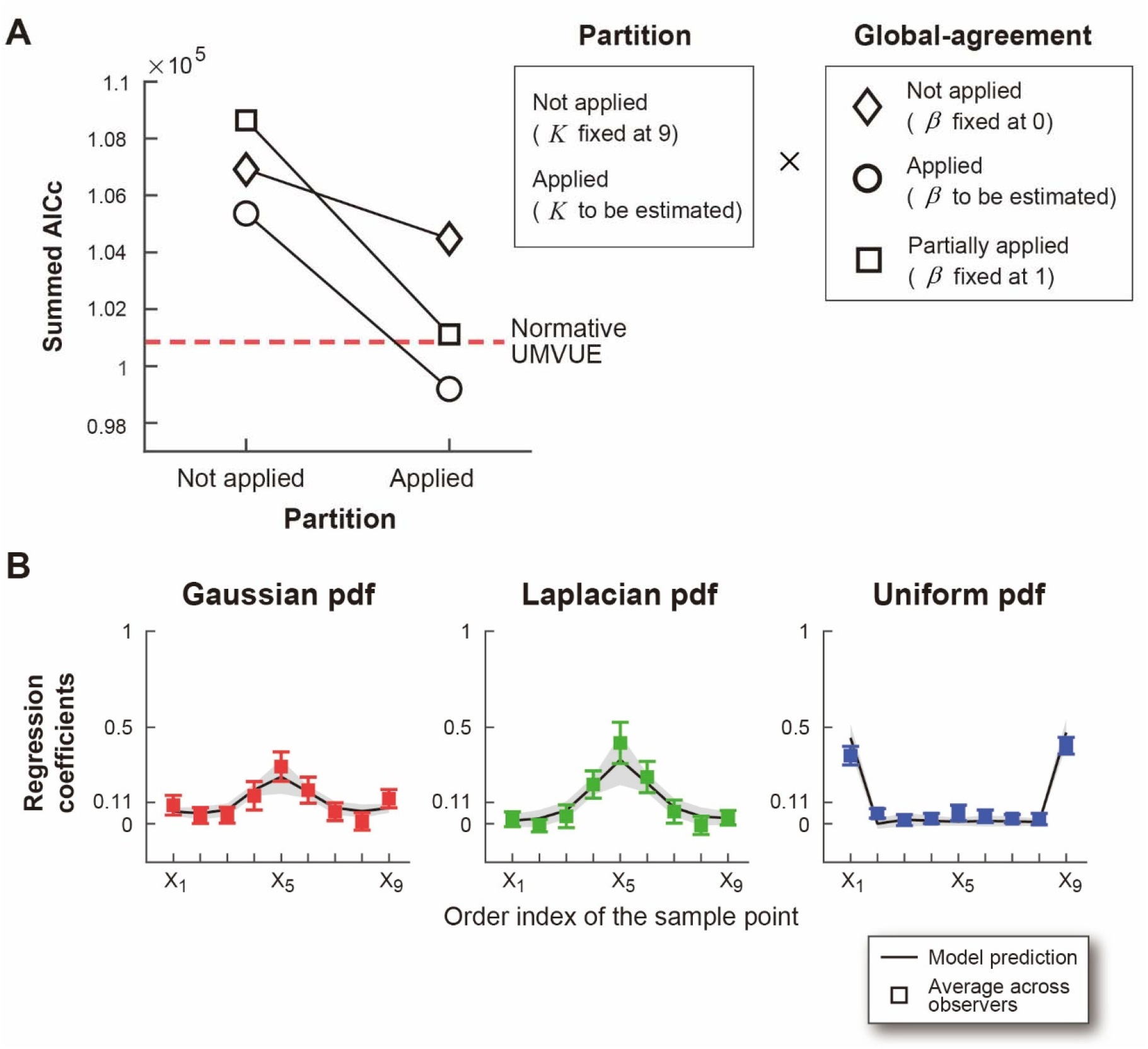
Visual Cluster Model Compared to Human Performance. **A. Factorial model comparison.** We built six models by combining two levels of partitioning (Step 1) and three levels of global agreement (Step 2). The y-axis denotes the AICc score summed across participants. The lower the AICc, the better the model. A dashed red line plots the AICc score for the normative UMVUE. A grouping parameter *K* is the number of clusters in K-means clustering we used to fit to the data. A parameter ***β*** is inverse temperature in the softmax transformation which converts the log-likelihood values to the relative weights (Figure 5). **B. Prediction of the Estimated Weights (Influence)**. Regression weights estimated from the data are taken from Figure 4 and replotted as colored squares. The model prediction of the estimated weights for each location task was estimated using a linear regression on the visual estimates recovered from the best Visual Cluster Model (Partition applied × Global agreement applied). The prediction of the estimated weights was averaged across observers for each task and was plotted as a black solid line. Shaded gray area denotes 99.7% confidence interval (± 3 SEM).

We estimated the influence of each point in the sample on the visual estimates generated by the two-stage VCM. The prediction of the influence measure by the model captured human data well in all three distributional families (Figure 6B). Specifically, the model produced a W-shaped influence pattern in the Gaussian task, a median-biased influence in the Laplacian task, and the average of extremes in the Uniform task. It is remarkable that a single model can generate such different estimators each tailored to a distribution family.

Patterns of predicted influence by other variants of the model are summarized in Supplementary Figure 5. When we applied the global agreement partially (***β*** fixed at 1) and had grouping parameters free, the model still produced distinct patterns of measured influence in the three location tasks (Supplementary Figure 5F), suggesting that we are not overfitting the data by including the global agreement parameter. However, without applying global agreement ***β*** (fixed at 0), we failed to properly fit the data (Supplementary Figure 5A&D). The models without partition (*K* fixed at 9) also failed to fit the data (Supplementary Figure 5A-C). These results again demonstrate that both partitioning and global agreement are essential computations.

Finally, we tested whether our results depend on the choice of the clustering algorithm. We repeated the model fit using a hierarchical clustering with single linkage algorithm (Nielsen, 2016; Shetty & Singh, 2021) and found that the results were almost identical (Supplementary Figure 6).

## Discussion

A central task of visual perceptual organization is to model how the visual system combines ‘parts’ into ‘wholes’ (Groen et al., 2017; Peirce, 2015; Ullman, 1984; Ullman et al., 2002). When parts have spatial attributes such as location, how does the visual system combine the locations of the parts into an estimate of the location of the whole? We considered a statistical version of this problem modeled on parametric point estimation (Lehmann, 1998). We asked observers to estimate the center of the pdfs of three different location families of one-dimensional pdfs. Observers knew everything about the pdf except for its location parameter which they were asked to estimate. In instructing the observers, we used the neutral term ‘center’ to refer to what was in fact the mean, median, and axis of symmetry of the pdf. The task is a paradigmatic statistical task: given a sample, estimate the parameters of the pdf that generated it. It is also a part of perceptual organization.

We compare human performance to that of a Uniformly Minimum Variance Unbiased Estimator (UMVUE) for that distribution. The UMVUE for each distribution is unbiased and has the lowest possible variance among location estimators. The normative minimum variance estimator is markedly different for the different distributional families we considered, namely the Gaussian, the Laplacian, and the Uniform.

How does the observer carry out the task? For the Gaussian location family, for example, the statistician needs only to compute the mean of the sample. The resulting estimate is the UMVUE but, as we have seen this is not what human observers do. We sought an alternative computation that would reproduce what human observers do, and that could be applied to estimation problems for the non-Gaussian distributions as well. *We propose that the visual system bases its estimates on the clusters in the sample and integrates the locations of the clusters, weighting them by how well each cluster alone accounts for the observed sample*.

Using influence measures, we tested whether observers adopt the same estimator for the three different distributions. We found that observers used different location estimators for the three distributions. We also tested whether the estimator for each distribution was the UMVUE for that distribution. The observers’ rules were different from UMVUE for Gaussian and Uniform location tasks but we could not reject the hypothesis that observers used the UMVUE in the Laplacian location task.

In the Gaussian location task, observers exhibit a systematic bias that deviates from equal weighting of each sample point, demonstrating a W-shaped influence pattern – placing large weights on the median and the two extreme points. This result is counterintuitive because it suggests that observers are not simply averaging visual elements, despite averaging being an elementary statistical operation that they are presumably familiar with. Deviations from equal weighting in the Gaussian location task have also been confirmed in previous studies, which show that observers’ estimates are biased by outliers in a sample (Juni et al., 2010; Moreland & Boynton, 2017; Rashid & Chubb, 2021). However, the influence patterns were flexibly adapted to the distributional constraints. In the Uniform location task, the two extreme sample points exerted strong influence on the observers’ estimates, while their influences were minimal in the Laplacian. Intuitively, the extreme points, if these are drawn from the Laplacian distribution, are poor indicators of the location of the Laplacian distribution. If observers were estimating the likelihood of the location of the distribution, the influence of these extreme points would be expected to be small in the Laplacian location task.

It is implausible that the human visual system has different rules of location estimation for any choice of location family. We wondered whether a common estimation algorithm, ideally based on Gestalt structure, could account for how the visual system assigns locations to samples from arbitrary location families.

We developed a two-stage Visual Cluster Model (VCM). Our model comparison analysis supported the two stages on the VCM. The first partition stage edits individual visual points so that points can be grouped into a cluster which itself defines a location. At the second stage (global agreement), the VCM evaluates how well each cluster approximates the location of the distribution generating the points. A cluster with a high likelihood value is weighted more in the visual judgment than a cluster with a low likelihood value.

The VCM then combines the cluster locations using weights that are proportional to the likelihood values assigned to clusters. The distributional constraints provided information that was necessary to estimate the likelihood of the location parameter of the distribution (Eq. 9). We demonstrated that a two-stage VCM simultaneously accounts for distinct patterns of the observers’ visual judgments across all three location tasks. We note that the VCM is not an L-estimator as the location estimator by the VCM is not the weighted sum of the order statistics of a sample. However, the VCM produces good approximations of observers’ estimators and therefore provides a descriptive account for how their location estimations were produced.

Sun et al (2019) also used a clustering algorithm to model the representation of human visual experience. In their task, observers were asked to estimate the location of the empirical mean or mode of the sample of 70 visual points, and points were drawn from a mixture of distributions that had three different peaks. The nature of their stimuli likely enhanced the visual segmentation of points beneath each peak, making clusters more salient. In contrast, the cluster structure in our task may be visually less salient, as we generated just 9 points from a unimodal (Gaussian, Laplacian, or Uniform) distribution. Nevertheless, models without the partition stage did not explain the data well.

Gestalt psychologists (Koffka, 1935/2013; Wagemans, Elder, et al., 2012; Wagemans, Feldman, et al., 2012) identified Gestalt rules underlying perceptual organization (for review see (Kubovy & Gepshtein, 2003)). Visual objects are grouped together based on a similarity of spatial attributes and these new *Gestalten* are assigned attributes including locations in space. The theory is descriptive and provides no clear justification for why Gestalten such as clusters are part of our visual experience (Jakel et al., 2016). The VCM offers two possible reasons for this.

First, the visual system can reduce the amount of information that must be processed by aggregating parts (Stigler, 2016). This becomes progressively advantageous as the number of visual elements increases. It is not simply the clusters that are useful but the fact that there can be many fewer clusters than sample points. Second, a cluster of points triggers useful information for visual judgments beyond just spatial attributes. In the VCM, each cluster is assigned its own location, which serves as an anchor where the likelihood of the center of the generating pdf is evaluated. By combining these two advantages, the VCM reduces the number of computations required to carry out the task, analogous to the resource rational principle in perception, cognition and decision-making (Lieder & Griffiths, 2019). For instance, in Figure 5C, the likelihood of the location of the pdf underlying the sample points is evaluated only at five cluster locations. Nevertheless, it approximates the maximum likelihood estimation as if we applied a low-pass filter at the limited sampling points. The resulting estimated likelihood function is not perfectly accurate. To compensate for this, the VCM computes a weighted average of these sampling points based on their estimated likelihoods. In this sense, Gestalten are not just fascinating visual illusions but may also correspond to intermediate steps in optimal computations.

In Marr’s framework of three levels of analysis (Marr, 1982), the VCM primarily addresses the computational and algorithmic levels of analysis, concerning how the visual system assigns a location to a ‘whole’ composed of ‘parts’ and what objective this computation may serve. The goal of the VCM can be interpreted as balancing accurate estimation against computational costs by replacing individual parts with a smaller number of perceptual clusters and evaluating the global agreement of these clusters with respect to the location of the whole. This interpretation is consistent with resource rational accounts of perception and cognition (Griffiths et al., 2015) in which the optimal computations are constrained by limited computational resources. It is also compatible with Bayesian inference in which the visual system approximates the most plausible interpretation of the structure of the image given its constituent visual elements (Froyen et al., 2015; Gershman et al., 2016; Shivkumar et al., 2025).

The VCM also provides a starting point for considering the algorithmic implementation of this computation. In this study, we instantiated the partition stage using two standard clustering algorithms: centroid-based K-means clustering (Hartigan & Wong, 1979) and connectivity-based hierarchical clustering (Nielsen, 2016; Shetty & Singh, 2021). We found that both algorithms reproduced behavioral patterns equally well, suggesting that the critical computation may be the formation of a cluster representation rather than a particular clustering algorithm. Recent work has likewise modeled human perceptual experience from a clustering perspective in tasks such as auditory streaming (Larigaldie et al., 2025), motion perception (Gershman et al., 2016; Shivkumar et al., 2025), and the representation of uncertainty (Sun et al., 2019). These studies employed a stochastic clustering approach called the Chinese restaurant process, assuming that perceptual noise leads to variability in the number of clusters. Such an approach may produce similar behavioral consequences to those generated by the clustering algorithms we used.

In conclusion, we addressed a central question in perceptual organization concerning how the ‘whole’ and its attributes are estimated from the attributes of individual ‘parts’. We formalized a statistical version of this problem by presenting observers with samples of points (‘parts’) drawn from three probability density functions (Gaussian, Laplacian, and Uniform) and asking them to estimate the location of the center of the pdf (‘whole’). We demonstrated that the observers’ visual judgments can be explained by a two-stage computational process: partition and global agreement. These results suggest that the observers’ estimate of the location of the ‘whole’ is based not directly on the local locations of the ‘parts’ but rather on the locations of ‘clusters’ they form. As such our findings imply that to further understand how the human visual system estimates the location of the ‘whole’, it is necessary to consider both the perceptual process which structures the groups based on ‘parts’ and the probabilistic structure (i.e., probability density function) that generates local locations of the ‘parts’.

## Materials and Methods

### Participants

Twenty participants (mean age 22.9, range 18–29, 13 females, 7 males, 0 other) participated in the experiment. The sample size was determined based on the previous location estimation studies employing comparable methodologies; eight studies included fewer than 10 participants per experiment (Gan et al., 2023; Juni et al., 2010; McGowan et al., 1998; Melcher & Kowler, 1999; Oruç et al., 2003; Rashid & Chubb, 2022; Sun et al., 2016, 2018) while one study included 16 participants (Sun et al., 2019). On this basis, a sample size of 20 participants was selected to ensure adequate statistical power. This study was approved by the University Committee on Activities Involving Human Subjects of New York University (IRB-FY2019-3450) and carried out in accordance with its approved guidelines. Written informed consent was obtained from each participant before the experiment. None of the participants was aware of the hypotheses under test in the experiment. Participants knew that they would receive $12 per hour plus a performance-related bonus (average bonus: $16.90, range: $11–$24). The experiment took approximately 2 hours.

### Apparatus

Stimuli were displayed on a horizontal monitor (VPIXX, VIEWPIXX, 514 mm × 288 mm). The monitor resolution was 1920 × 1080 pixels with a 60-Hz refresh rate and we refer to location on the monitor by horizontal coordinates ranging from −257 mm to 257 mm and vertical coordinates −144 mm to 144 mm. The participants were seated at a viewing distance of approximately 43 cm and manipulated a computer mouse to carry out tasks. All stimuli were presented and controlled using the Matlab Psychophysics Toolbox (Brainard, 1997; Pelli, 1997).

### Training Task

Participants were first familiarized with the three distributional families: Gaussian, Laplacian, and Uniform (see Supplementary Figure 1A). In this training stage, participants were not yet asked to make any estimates but were asked to observe the population pdfs, the population means, and the samples drawn from these population distributions. On each trial, we plotted a pdf drawn from one of the three distributional families and a sample from the same pdf. The origin (0,0) was set at the vertical and horizontal center of the screen. The location ***μ*** of the pdf was drawn uniformly from the interval −100 mm and 100 mm. Following this, a sample of *N* points was randomly drawn. The sample points were displayed underneath the shape of the pdf. The points were discs with a radius of 1.5 mm and colored by the same color that was used to draw the underlying pdf (for instance, red Gaussian; green Laplacian; blue Uniform). The color codes were counter-balanced across participants.

The population variances ***σ***^**2**^ of all pdfs were fixed at 2500 mm^2^ (σ = 50 mm). We added vertical jitter to the vertical positions of the sample points to reduce the chance that sample points would occlude one another. The variance of vertical jitter was fixed at 6.25 mm^2^. To aid the observer in interpreting the pdf and sample during training, we marked the location ***μ*** of the pdf by a white solid line segment and dashed lines (width: 2 mm, height: 35 mm).

During training, the sample size *N* was initially 300 and was gradually reduced to 50, 20 and finally 9. With 300 points the displayed sample could be readily matched to the pdf from which it was drawn.

Presentation of the sample when N > 9 was extended across time: 18 additional points were drawn every 100 ms from the underlying pdf until the required number of points (300, 50, 20, or 9) points appeared on the screen. The sample points were displayed for 2s. After an inter-trial interval of 1s, a new pdf with randomly chosen location ***μ*** was displayed together with a sample drawn from the pdf.

A distribution was chosen from one of the three distributional families (Gaussian, Laplacian, and Uniform) and was repeated for five trials. In each trial, a new sample was generated from the distribution chosen. After 15 trials with the presentation of 300 sample points (five trials for each of the three distribution families), the second block with 50 sample points started. The third block and the fourth block used 20 and 9 points, respectively, giving 60 trials in total. The final sample size (*N =* 9) was the one used in the main experimental task, the location task. The training task provided substantial opportunity to observe typical Gaussian, Laplacian, and Uniform samples while learning the color codes.

In the instructions, we used the following analogy:

On every trial, you will first see a white apple tree located somewhere on the screen. You will then see a curve which tells you where the apples are most likely to fall. The higher the curve is, the more likely an apple will fall below that point on the curve. Where the curve is high, there probably will be more apples. Where the curve is low, fewer. The density curve is always symmetric around the apple tree. The horizontal location of the apple tree is not fixed but it will vary from trial to trial along with the curve and the apples. The tree, the apples and the curve will all move together from left to right on each trial. There are three types of density curves from where apples will fall, each coded by color. One is bell-shaped (red), the second is “pointy” (green), and the last is like a box (blue).

### Visual Location Task [Main Experiment]

The stimuli in the visual location task were the same as in the visual training task with the following differences: On each trial in the Visual Location Task, participants saw only a color-coded sample of N = 9 points randomly drawn from the Gaussian, Laplacian or Uniform distribution. The pdf and the location of the pdf were not shown on the screen. To help participants remember the three density curves, small icons representing the shape of the three distributions were drawn near the top of the screen (see Supplementary Figure 1B and Supplementary Videos 2&3).

Participants were asked to estimate the unknown position of the center (i.e., location ***μ***) of the underlying pdf using only the positions of sample points. They could move a white upward pointing arrow (10 mm) to any location on the screen using a mouse. Participants set the arrow at a position where they believed the horizontal center of the underlying pdf was located and clicked the mouse button to record their estimates. We ignored the vertical displacement of the arrow in analyzing the observers’ settings.

Following a response, we presented a white vertical line (35 mm) marking the location of the underlying pdf as feedback on the error between the participant’s estimates and the true position of the center of the pdf. Participants received 500 – *D*^*2*^ points where *D*^*2*^ is the squared magnitude of their error in mm. If the position of the estimates perfectly matched the true center (*D*^*2*^ = 0), the participant received 500 points. Points received could be negative if the absolute estimation error exceeded 22.4 mm. At the end of each trial, the estimation error (mm), the score (points) and the cumulative score were presented as a graphical display for 1.5 s. Participants received a bonus payment proportional to the total score they earned across all trials ($1 per 10,000 points). The estimator that maximized the participants’ expected bonus for each distribution family was the UMVUE for that distribution family.

We provided warm-up trials before the main experiment. During warm-up, the sample size was initially 300 and gradually reduced to 50, 20, and 9. The warm-up task became more difficult as the sample size was reduced. As in the training task, a distribution was chosen from one of the three distributional families (Gaussian, Laplacian, and Uniform) and was repeated for five trials. In each trial, a new sample was generated from the distribution chosen, giving 15 = 5 x 3 trials for the first block of 300 sample points. Then the second block of 50 sample points started. The third block and the fourth block used 20 and 9 points, respectively, giving 60 warm-up trials in total. Following a short break, the main experiment started. One block consisted of 10 trials. Samples from the three distributions were interleaved across blocks. Each participant ran 90 blocks (30 blocks for each distribution) with a short break after every 6 blocks.

### Uniformly Minimum Variance Unbiased Estimator among L-estimators

The Laplacian location family has no closed-form solution for its UMVUE. Therefore we computed the unbiased L-estimator that minimized variance numerically as follows. We denote the UMVUE among L-estimators as *UMVUE*_*L*_.

Given the n × n covariance matrix Σ of the order statistics ***X***_**1**_, … ***X***_***N***_ we can compute the choice of weights that leads to a minimum variance estimator as

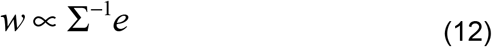

where *e* = [1,1,…,1]′, assuming that the correlations between the random variables are non-zero (for proof, see Appendix A in Oruc et al., 2003), which is our case because the random variables are ordered. We ran the numerical simulation by resampling the order statistics 1,000,000 times and then computing the inverse of the covariance matrix and the weights in Eq. 12. We normalized the weights so they sum to 1.

### Measuring influence

*Order statistics*. We performed a regression analysis to estimate the influence of each point in the sample on the participant’s estimate of the position of the horizontal location of the distribution (Gaussian, Laplacian, or Uniform) that had generated the sample. We ordered 9 sample points in each sample in ascending order from the leftmost point to the rightmost point. For each of the three distributions, we regressed the vector of the participant’s estimators **T**^**o**^ which contained 300 data points by the 300 by 9 matrix **X**. Columns were vectors of order statistics for each trial ***X***_**(9)**_, … ***X***_**(9)**_. Our linear regression model is thus 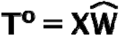 where 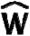 is the estimated weight vector. We used an ordinary least squares regression with the constraint that weights add to 1: 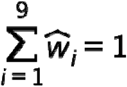. We estimated influence for each participant and each distribution family of the population distribution. In addition, we estimated the weight vector of the visual cluster model by regressing the vector of the visual estimates predicted by the VCM by the matrix. For each variant of the VCM, we retrieved their visual estimates from the best fit parameters in each model (see *Model fitting and evaluation*). See Supplementary Figure 5 for the estimated weights in each model. The influence measures are a sensitive way to uncover how participants behave in carrying out the task and discover discrepancies between human performance and optimal performance (Dal Martello et al., 2023; Ota et al., 2024; Ota et al., 2026; Ota et al., 2025a; Ota & Maloney, 2024).

### Model fitting and evaluation

For each participant, we fitted the model’s visual estimates (Eq. 11) to the participant’s estimates by minimizing the negative log-likelihood of the observed estimates using Bayesian adaptive direct search (BADS) (Acerbi & Ma, 2017). We assumed that the observers’ location estimate T^O^ ~ N(T^VCM^, c^2^) was Gaussian distributed.

In the first stage of the VCM, we tested two levels of partitioning sample points. At the first level, we did not apply partition and fixed the grouping parameter *K* (i.e., the number of clusters) in K-means clustering at 9 across all three location tasks. At the second level of applying partition, we estimated *K* separately from data in each of the three location tasks as a free parameter. Because *K* must be an integer, we estimated *K* by a grid search in steps of 1. In each iteration of the grid search, the same integer value of *K* was used across trials in each task. The estimated value of *K* could differ across the three location tasks as the tasks for the three conditions were interleaved. We optimized other parameters (described below) for each combination of the values of the three grouping parameters *K*_*G*_, *K*_*L*_, *K*_*U*_ and chose the best combination that minimized negative log-likelihood on each individual’s data.

In the second stage of the VCM, we tested three levels of global agreement. At the first level, we did not apply global agreement and fixed the parameter *β* (the inverse temperature in the softmax decision rule) at 0. At the second level of applying global agreement, we treated *β* as a free parameter with the boundary *β* ∈[0,4]. Here we fitted the value of *β* jointly across the three location tasks. Because the probability density did not convey any information about how the relative weights should be assigned to each cluster, we reasoned that participants did not adjust the global agreement parameter according to the distribution family. Finally, at the third level, we partially applied global agreement by fixing *β* at 1. Therefore, this factorial design allowed us to compare six variants of the model. In all six models, we included a Gaussian response variance σ ^2^ and a constant bias parameter *b* for individual’s location estimates with the boundary *b* ∈ [−50,50]. Again, we fitted the values of σ ^2^ and *b* jointly across the three location tasks.

We also used hierarchical clustering with single linkage to test whether the results of the VCM depend on a clustering algorithm we chose. Instead of *K*, we used 0 E [0,150] as a grouping parameter. We fitted *θ*_*G*_, *θ*_*L*_, *θ*_*U*_ to the data while other parameters remained the same (Supplementary Figure 6). This parameter is a cutoff threshold in millimeter. If the distance between two sample points or clusters is below a threshold, these two are grouped into one cluster.

In the normative model, we fitted UMVUEs matched to the distribution family to the data in the corresponding location task with two free parameters σ ^2^ and *b*. In all models, we repeated the search process with different starting points to verify that we had found the global minimum. For model comparison, we applied AICc—Akaike information criterion with a correction for finite sample size—to each participant and each model as the information criterion for goodness-of-fit (Burnham & Anderson, 2002; Hurvich & Tsai, 1989). The Bayesian information criterion (BIC) is motivated as a dimension-consistent model selection criterion designed to select the “true model” when it is contained in the candidate models being considered. Burnham & Anderson (2002) argues that expecting any candidate model to represent the “full truth” is often unrealistic in biological sciences. They therefore recommend AIC-type criteria for the analysis of empirical data, because AIC is not designed to identify the true model but rather to select the model expected to be closest to the data-generating process by minimizing the relative information loss among the candidate models. The AIC’s justification is asymptotic and it can be biased toward more complex models when sample size is small relative to the number of model parameters. AICc introduces an additional penalty term that reduces this small-sample bias, helping to avoid potential overfitting and favour more parsimonious models. Based on AICc values across participants and across models, we computed the protected exceedance probabilities (Rigoux et al., 2014; Stephan et al., 2009) using a Matlab function ‘spm_BMS’ in SPM 12 (https://www.fil.ion.ucl.ac.uk/spm/). The sign of AICc values was reversed by multiplying with 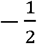 so that an entry to the function ‘spm_BMS’ can be converted to log model evidence (https://mbb-team.github.io/VBA-toolbox/wiki/BMS-for-group-studies/).

## Author’s Contributions

A, B, C, and D designed the experiment and analysis, A and B collected data. A carried out the analyses. All authors contributed to developing and evaluating models and writing the article.

## Data and code availability

The experiment programs used in this study, pre-processed data and analysis scripts are publicly available at https://osf.io/g7p63.

## Supporting Information

**Supplementary Figure 1.**
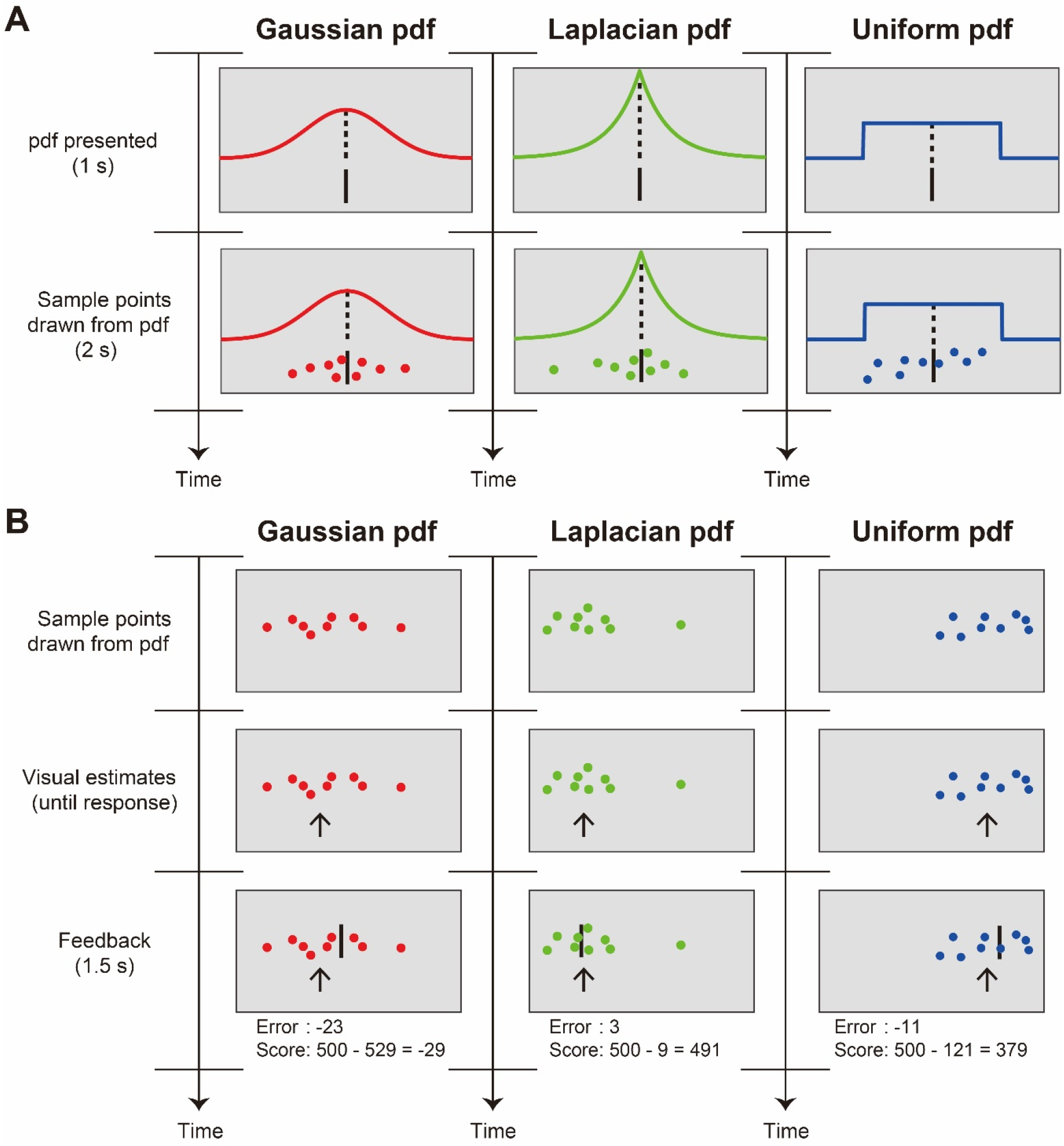
Training Task and Visual Location Task. **A. Training Task**. Three distributional families were color-coded. On each trial, we presented one of the three pdfs with varying its location ***μ*** along the horizontal axis and a sample drawn from the same pdf. The sample size reduced from *N* = 300 to *N* = 9 between blocks. Participants were asked to observe a color-coded sample drawn from the pdf with the same color code. **B. Visual Location Task**. On each trial, a sample of 9 points is drawn from one of the three distributional families with its location varied along the horizontal axis. Participants were asked to estimate the location of the invisible center of the pdf that had generated the sample by moving an arrow horizontally. Following a response, the true location of the center of the pdf was presented as a vertical line. A squared error and a score for a trial were also provided as feedback.

**Supplementary Figure 2.**
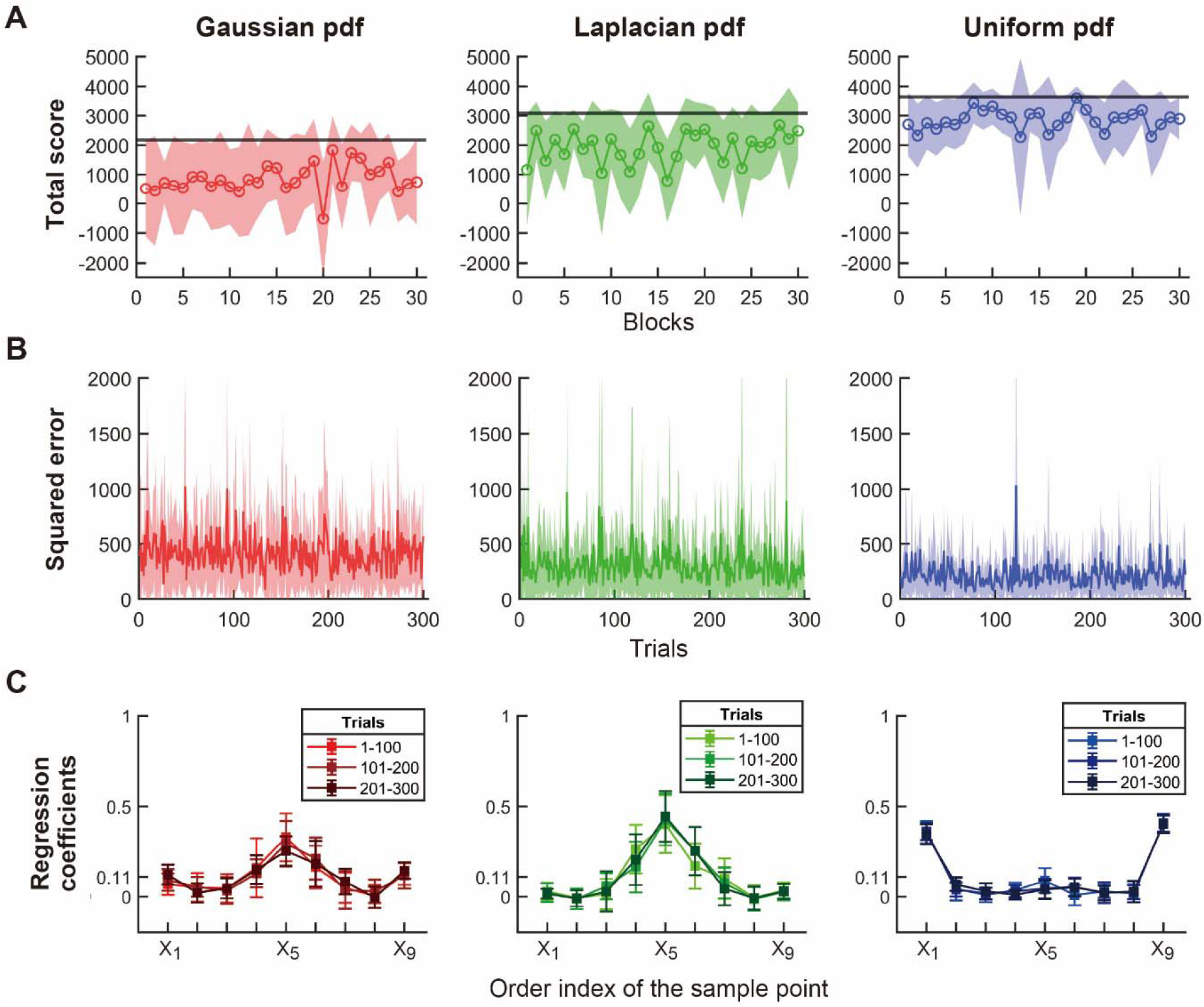
Behavioral Performance did not Change Systematically Across Trials. **A. Total Score**. The participant’s total score in each location task is plotted across blocks. Data are averaged across all participants. The shaded areas indicate 99.7% confidence interval (± 3 SEM). Black horizontal lines are the theoretically maximum total score obtained by UMVUE that corresponds to each distribution family. Two-way within participant ANOVA, using the number of blocks (30) and the distribution family (3) as independent variables, shows neither main effect of the block (*F* [29, 551] = 1.43, *p* = 0.069, partial *η*^2^ = 0.070) nor a significant interaction between the block and the distribution (*F* [58, 1102] = 1.19, *p* = 0.155, partial *η*^2^ = 0.059). **B. Squared Error**. The squared error between the participant’s visual estimate and the true location of the center of the distribution is plotted versus trial. Two-way within-participant ANOVA, using the number of trials (300) and the distribution family (3) as independent variables, shows neither main effect of the block (*F* [299, 5681] = 1.11, *p* = 0.095, partial *η*^2^ = 0.055) nor a significant interaction between the block and the distribution (*F* [598, 11362] = 1.08, *p* = 0.091, partial *η*^2^ = 0.054). **C. Estimated Weights (Influence)**. To additionally confirm a lack of learning effect over trials, we computed influence measures separately for the first, the second, and the last 100 trials using a linear regression. The estimated weights are averaged across all participants. The error bars indicate 99.7% confidence interval (± 3 SEM). A two-way within participant ANOVA, using the order of points (9) and the bin of trials (3) as independent variables, shows no significant interaction between the order index of the sample point and the trial bin in the Gaussian location task (*F* [16, 304] = 0.70, *p* = 0.796, partial *η*^2^ = 0.035), in the Laplacian location task (*F* [16, 304] = 0.74, *p* = 0.751, partial *η*^2^ = 0.038), nor in the Uniform location task (*F* [16, 304] = 1.22, *p* = 0.250, partial *η*^2^ = 0.060).

**Supplementary Figure 3.**
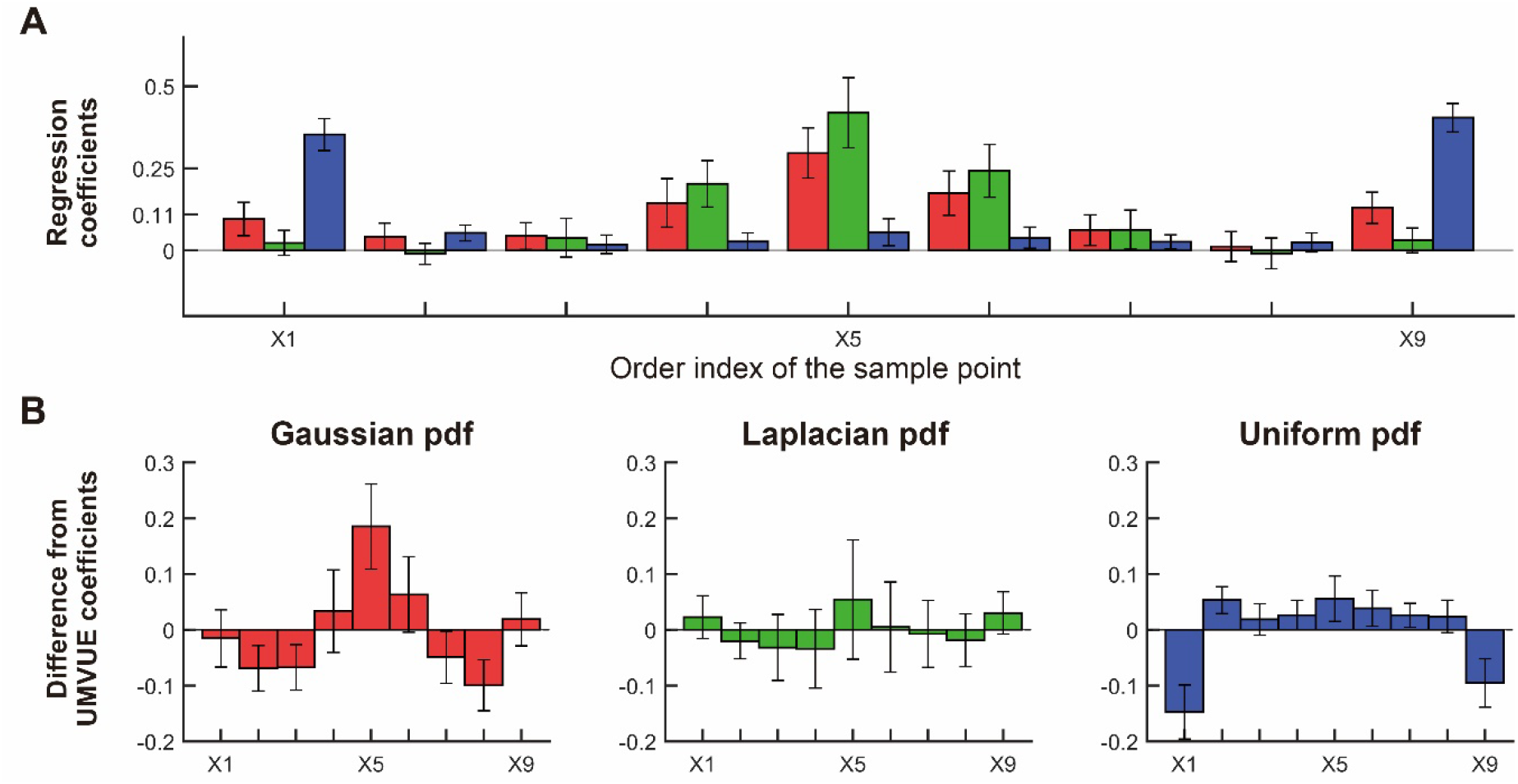
Pairwise Comparisons of the Estimated Weights. **A**. The values of the estimated weights (influence measure) from the data are taken from Figure 4 and replotted as a bar graph as a function of the order statistics. **B**. Difference between the estimated weights and the weights employed by the UMVUE (for the Laplacian the UMVUE_L_). In all panels, bars represent the average across participants. The error bars indicate 99.7% confidence interval (± 3 SEM).

**Supplementary Figure 4.**
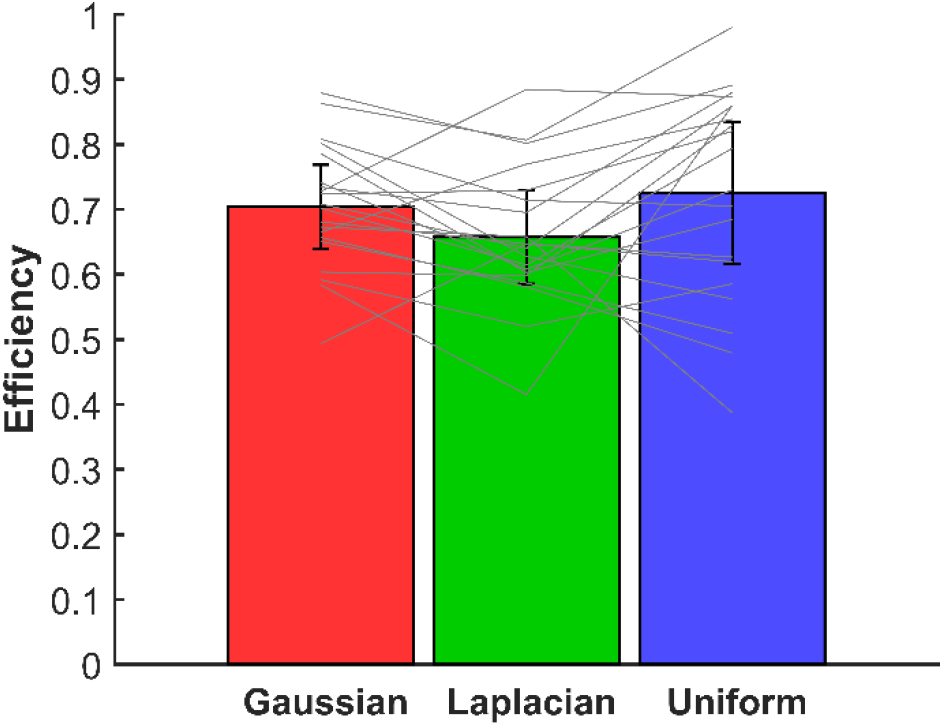
Efficiency of the obervers’ estimators. For each of the three location tasks and for each observer, we computed the variance of observers’ estimator across trials and the variance of the UMVUE (which matched each location family) across trials. We then calculated the efficiency based on Equation 4. Bars represent the average across participants, whereas gray lines represent individual participants. The error bars indicate 99.7% confidence interval (± 3 SEM).

**Supplementary Figure 5.**
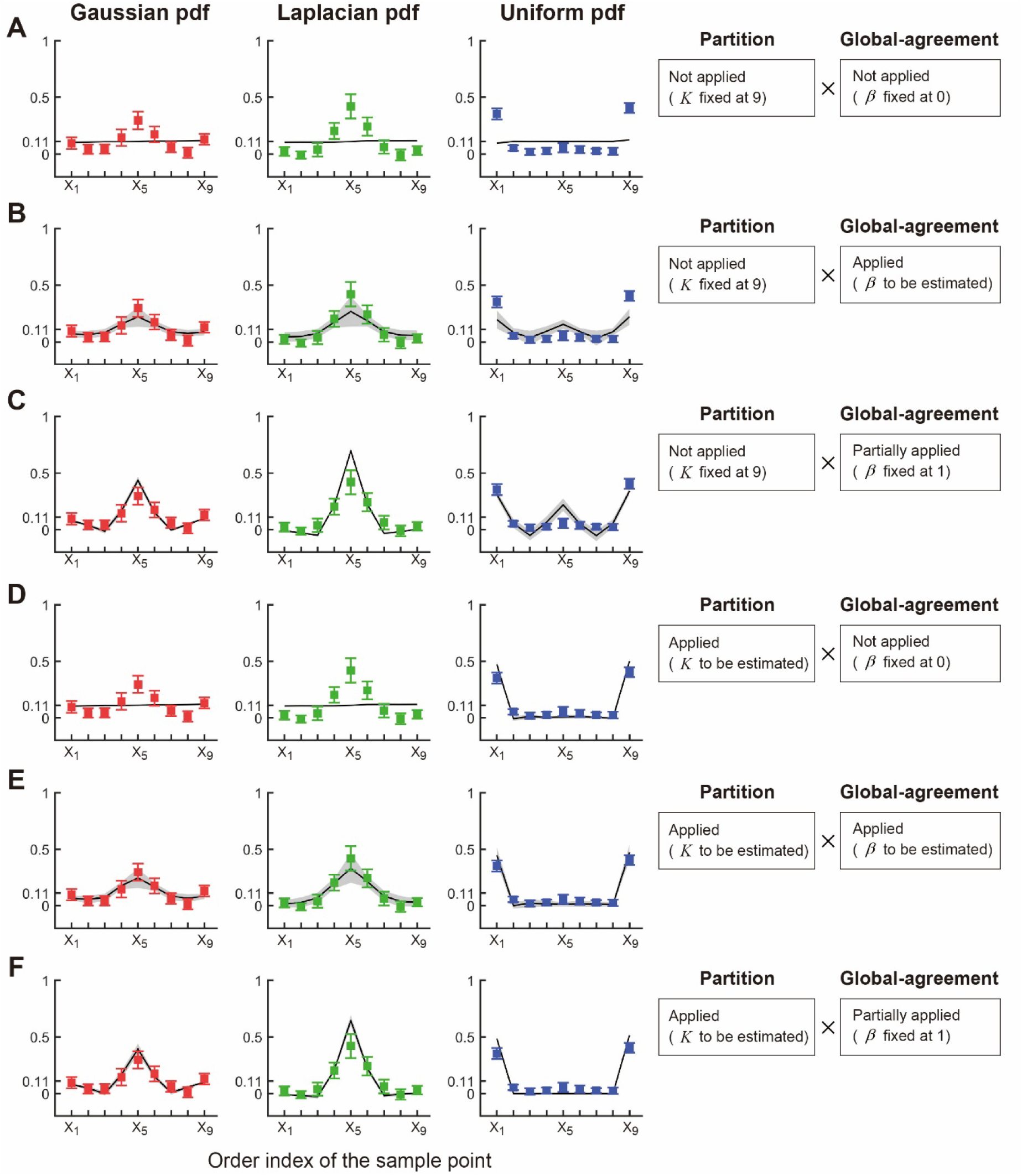
All Variants of the Visual Cluster Model. The predictions of the estimated weights for each model in the factorial model comparison. Regression weights estimated from the data are taken from Figure 4 and replotted as colored squares. Mean model fits across observers are plotted as black solid lines while shaded gray areas denote 99.7% confidence interval (± 3 SEM).

**Supplementary Figure 6.**
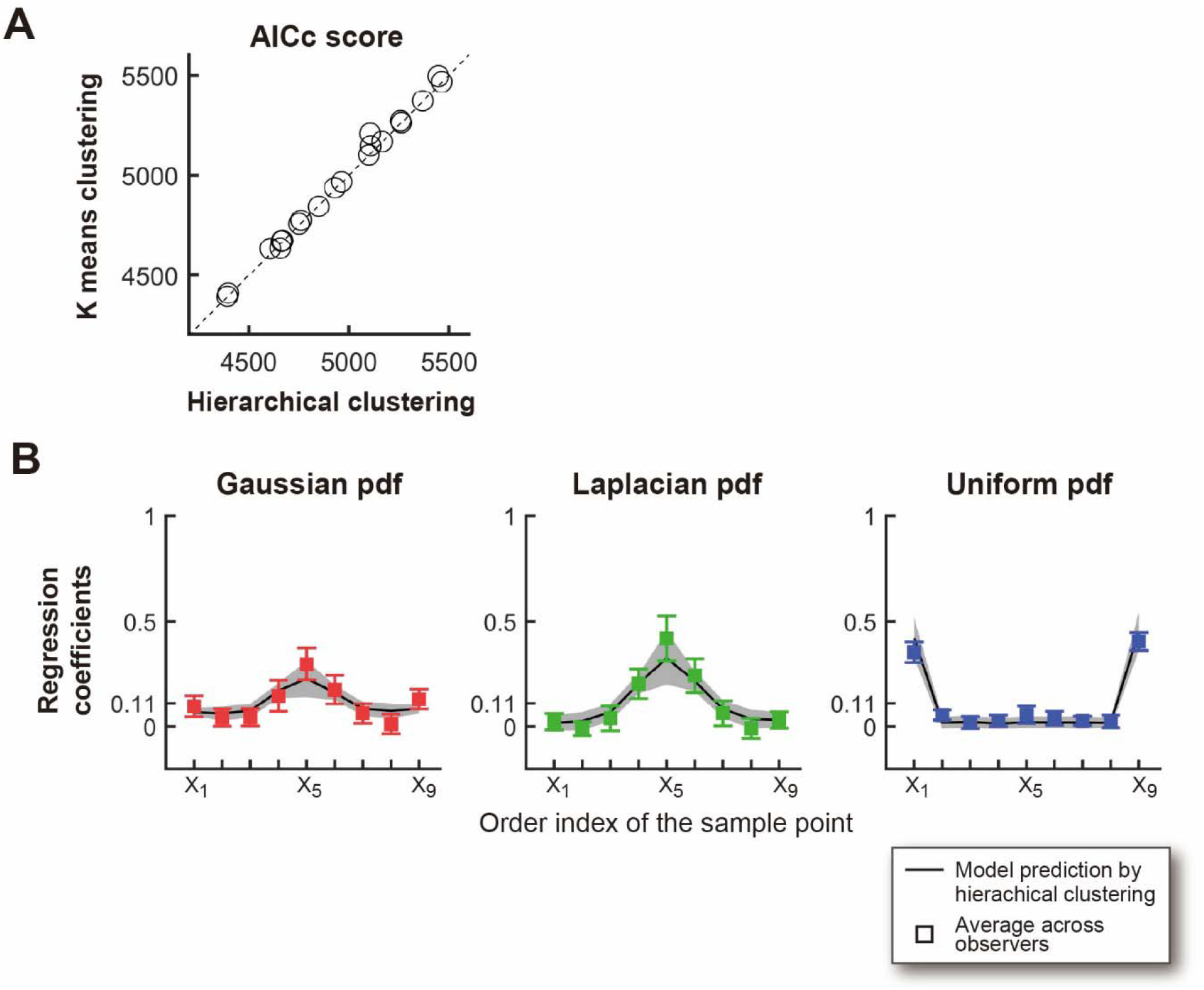
Comparison of the Model Fit across Two Clustering Algorithms. **A**. A Comparison of AICc Scores. We adopted K-means clustering in the main text to fit the visual cluster model to the data. We repeated the clustering algorithm analysis using hierarchical clustering with single linkage. **B**. Prediction of the Estimated Weights (Influence). Regression weights estimated from the predictions of the VCM using hierarchical clustering. AICc scores and the estimated weights are almost identical between the two clustering algorithms (see Figure 6).

## Footnotes

1 The actual PDFs used were always scaled to have the same variance. We present them in simplified form in Eqs. 2 and 3. See Methods.

